# ‘Chromatic’ Neuronal Jamming in a Primitive Brain

**DOI:** 10.1101/496745

**Authors:** Margarita Khariton, Xian Kong, Jian Qin, Bo Wang

## Abstract

Jamming models developed in inanimate matter have been widely used to describe cell packing in tissues^1–7^, but predominantly neglect cell diversity, despite its prevalence in biology. Most tissues, animal brains in particular, comprise a mix of many cell types, with mounting evidence suggesting that neurons can recognize their respective types as they organize in space^8–11^. How cell diversity revises the current jamming paradigm is unknown. Here, in the brain of the flatworm planarian *Schmidtea mediterranea*, which exhibits remarkable tissue plasticity within a simple, quantifiable nervous system^12–16^, we identify a distinct packing state, ‘chromatic’ jamming. Combining experiments with computational modeling, we show that neurons of distinct types form independent percolating networks barring any physical contact. This jammed state emerges as cell packing configurations become constrained by cell type-specific interactions and therefore may extend to describe cell packing in similarly complex tissues composed of multiple cell types.

---

Jamming behaviors have been extensively studied in inanimate systems undergoing liquid-to-solid transitions, including foams, gels, and emulsions^17–21^, and extended to describe cellular packing in biological tissues^1–7^. In physical systems, jamming transitions are controlled by a set of canonical variables: density, temperature, and stress^18,20,21^. In tissues, these variables have been linked to cellular properties, including shape, migration, and mechanics^4,6^. However, cell diversity has not been accounted for in the current framework of jamming, despite complex tissues being comprised of many distinct cell types. In particular, neural tissues possess extremely high cell diversity and it is well known that neurons can recognize and interact with their respective types as they organize in space^8,9,22^. A neuron can avoid other neurons of its same type even when they are several cell body sizes apart, mediated by a variety of broadly conserved biological mechanisms (**Fig. 1a** and **Supplementary Note 1**)^10,11,22^. This repulsion causes homotypic neurons to organize into two-dimensional arrays with regular inter-neuronal spacing in sensory systems^10,11,22,23^. How these type-specific interactions organize cells in more complex three-dimensional tissues is unknown.

**Figure 1.**
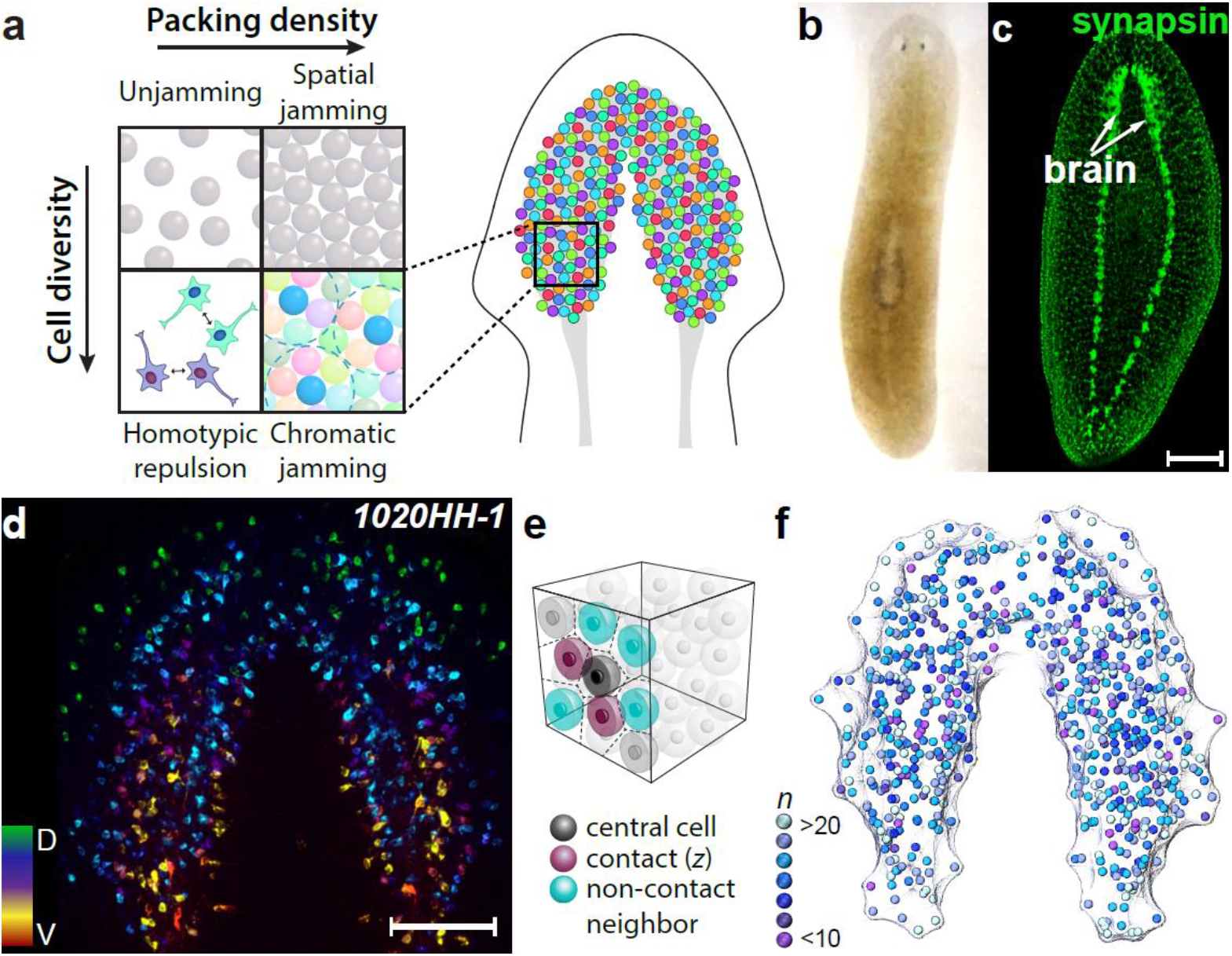
The planarian brain allows quantification of neuronal packing with single-neuron resolution. **a**) Diagram illustrating particle packing of a system as a function of packing density and cell diversity: low density and diversity gives a fluid, unjammed system (top left), while increasing the packing density in homogeneous tissues results in the conventional spatially jammed state (top right). Homotypic repulsion (bottom left) gives rise to a chromatically jammed state (bottom right), as in the spatial organization in neural tissues composed of numerous distinct neuronal types. Dashed circles: range of cell type-specific interactions. **b**) Brightfield image showing the gross morphology of the planarian, *S. mediterranea*. **c**) Fluorescence image (maximum intensity projections of confocal stack) depicting the planarian nervous system as stained for synapsin (using antibody 3C11, Developmental Studies Hybridoma Bank). Scale bar: 200 μm. **d**) Depth-colored projection image showing RNA-FISH of neuropeptide *1020HH-1* to reveal the spatial distribution of *1020HH-1*^+^ peptidergic neurons in the planarian brain. Colormap: dorsal-to-ventral. Scale bar: 100 μm. **e**) Illustrative diagram of homotypic neighbors and contacts. For a given cell (black), its homotypic nearest neighbors (blue and purple) are defined by adjacent Voronoi tessellation units, and neighbors in geometric contact (purple) are defined as units with touching enclosing spheres. Solid spheres: cell body centroids; translucent zones: range of homotypic repulsion. **f**) Centroids of *1020HH-1*^+^ peptidergic neurons determined from the image in **d**. Colors represent number of nearest homotypic neighbors (*n*) of each individual neuron, measured by Voronoi tessellation.

To address this question, we propose a new jamming transition, which is controlled by the cell diversity of a tissue and thereby represents an orthogonal dimension to previously characterized jamming behaviors. In previous studies, the jamming state manifests as the positions and shapes of cells are constrained by their physical neighbors^3–7^. Here, cell type adds an additional degree of freedom, which can be progressively reduced as the cell diversity of a tissue decreases (**Fig. 1a**). At a critical number of cell types, homotypic repulsion becomes so abundant as to propagate throughout the system and solidify cellular packing, analogous to gelation in random networks^24^. Inspired by the color map problem^25^, we term this transition ‘chromatic’ jamming.

Here we demonstrate chromatic jamming through experimental and computational model systems. Experimentally, we use quantitative 3D imaging to investigate neuronal packing within the brain of the planarian *Schmidtea mediterranea*, one of the most basal organisms known to have a demarcated central nervous system, which contains several dozen neuronal types^13,26,27^ (**Fig. 1c-d**). Importantly, the total number of neurons in the planarian brain scales linearly with animal size and can vary by more than one order of magnitude, from thousands to tens of thousands, while the number density of each neuronal type remains nearly constant^14^. This striking invariance suggests that neurons pack into a tissue size-independent, scale-free structure.

With the high genetic similarity to the vertebrate nervous system^13,15,26^, the planarian brain also provides the simplicity necessary for precisely determining individual neuronal positions. We acquired 3D confocal images of non-overlapping neuronal types using RNA fluorescence *in situ* hybridization (FISH) against a panel of neuropeptide genes, each specifically labeling the cell bodies of a distinct type of peptidergic neuron^13,14,28^ (**Fig. 1d** and **Supplementary Fig. 1**). Our imaging resolution of 350 nm lateral, 700 nm axial, is sufficient to resolve single cells with an average diameter of 6 μm using automated imaging analysis.

Two essential features are evident regarding the type-specific packing organization: First, despite the dense packing of neurons throughout the brain space, homotypic neurons are rarely in direct physical contact, consistent with the property of homotypic repulsion^9^; they do not organize into regular mosaic patterns either, unlike the well-studied 2D sensory systems^9,11,23^. Second, in alignment with previous reports in other animals (e.g., fruit fly and mouse)^9,10,22^, no interaction is observed between heterotypic neurons (**Supplementary Fig. 2**), allowing us to treat individual neuronal types separately.

To quantify the packing structure within each neuronal type, we located the centroids of individual neuronal cell bodies and identified their nearest homotypic neighbors using Voronoi tessellation (**Fig. 1e**). **Fig. 1f** plots the centroids colored according to the number of their homotypic nearest neighbors, n, with more examples shown in **Supplementary Fig. 1.** Although large variations in *n* between individual cells are observed, no long-range spatial inhomogeneity is apparent. **Fig. 2a** shows that the average number of nearest neighbors, 〈n〉 = 14 ± 0.2, is stable over various neuronal types and across a wide range of brain sizes. This value is in good agreement with that of classic 3D random packings in particulate matter^17–20^.

**Figure 2.**
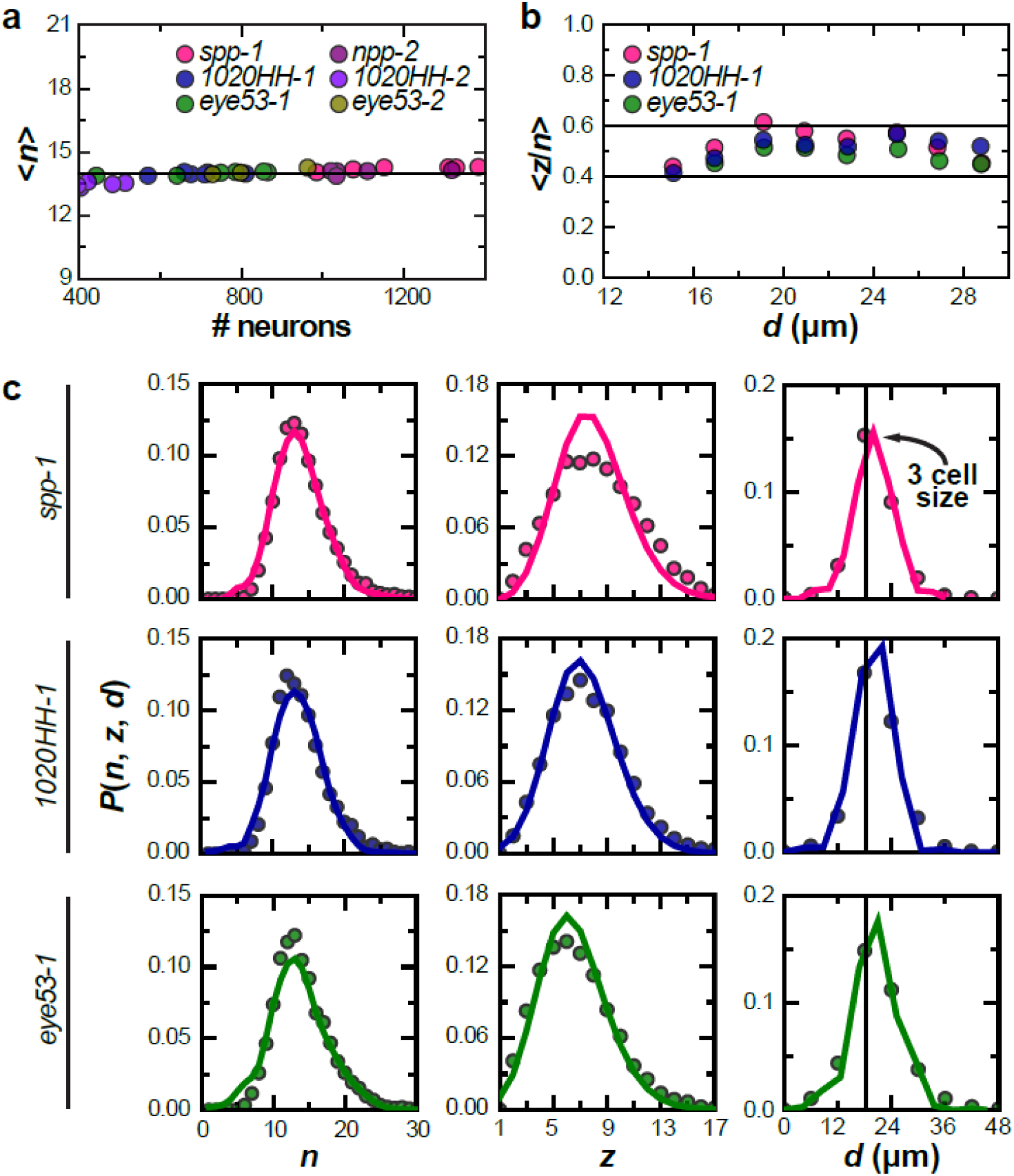
Neuronal packing exhibits geometric hallmarks of jamming. **a**) Average number of nearest homotypic neighbors, 〈*n*〉, plotted against total number of neurons in respective populations, is invariant for all 6 measured peptidergic neuronal populations from 34 planarians of various sizes. Horizontal line: 〈*n*〉 = 14. **b**) Average fraction of contact neighbors, 〈*z/n*〉, plotted against packing unit size, falls in the range of 0.4 to 0.6 (horizontal lines) regardless of packing unit size and neuronal type. **c**) Probability distributions of number of nearest neighbors (*n*, left), number of contacts (*z*, center), and packing unit size (*d*, right) for 3 representative peptidergic neuronal populations. Distributions of *n* and *z* are centered about 14 and 6.5, respectively, with the *d* distribution having a peak at 18 μm, equivalent to 3 cell sizes. Note that *P*(*n*) spans the range from 5 to 25 and stands in contrast with the narrow distribution (12-17) expected from the packing of monodisperse units^17^. Symbols: experimental data; solid lines: granocentric model of jammed polydisperse soft particles^17,19^.

If neurons truly pack into a jammed state with respect to their homotypic neighbors, then only a fraction of nearest homotypic neighbors should be in geometric contact^17,18^ (**Fig. 1e**). This fraction must be larger than the characteristic value of random loose packing (the loosest way to pack particles) and its average should be independent of local variables such as packing unit size^17–19^. To test these predictions, we fit the largest possible sphere to each Voronoi unit, defining packing unit size (*d*) as the diameter of the fit sphere, and units with touching spheres as geometric contacts. The average unit size, 〈*d*〉, is approximately 18 μm, corresponding to 3 cell sizes, consistent with the expectation that neurons prohibit direct contact of homotypic neighbors^9-11,22,23^. The average fraction of contact neighbors, 〈*z/n*〉, ranges from 0.4 to 0.6 (**Fig. 2b**), which is above the reported values of random loose packing (0.3-0.4) yet is consistent with random close packing near the jamming transition in non-living particulate systems^17–19^. As predicted, 〈*z/n*〉 is also largely independent of neuronal type and Voronoi unit size.

Consistently, the fluctuations of these geometric measures in the planarian brain also exhibit hallmarks of the jamming state. We chose the granocentric jamming model^17,19^ for quantitative comparison, as it provides closed-form expressions for the microscopic distributions of nearest neighbors and contacts. We measured the distributions *P*(*n*), *P*(*z*), and *P*(*d*) for three representative neuronal populations. **Fig. 2c** shows that theory and experiment agree quantitatively for all distributions, without any free fitting parameter. Minor discrepancies likely reflect the complex shapes of neurons.

These geometric measures collectively establish that homotypic interactions can determine the state at which neurons pack. This is a key feature of how chromatic jamming contrasts with conventional jamming, which is mediated by physical contacts. To understand how homotypic contacts build up to the chromatic jamming state, we constructed a minimum model to account for both physical and homotypic contacts. We modeled neurons as particles partitioned evenly into *c* types, with each type assigned a random representative color (**Fig. 3a**). The range of repulsion between heterotypic particle pairs is one particle size and that between homotypic pairs is three particle sizes, to match the experimentally measured average distances between heterotypic and homotypic neurons (**Fig. 2c** and **Supplementary Fig. 2**). Initial system configurations were generated randomly at a total packing density *ϕ* and equilibrated by shuffling particle positions to minimize total energy (**Supplementary Fig. 3a**).

**Figure 3.**
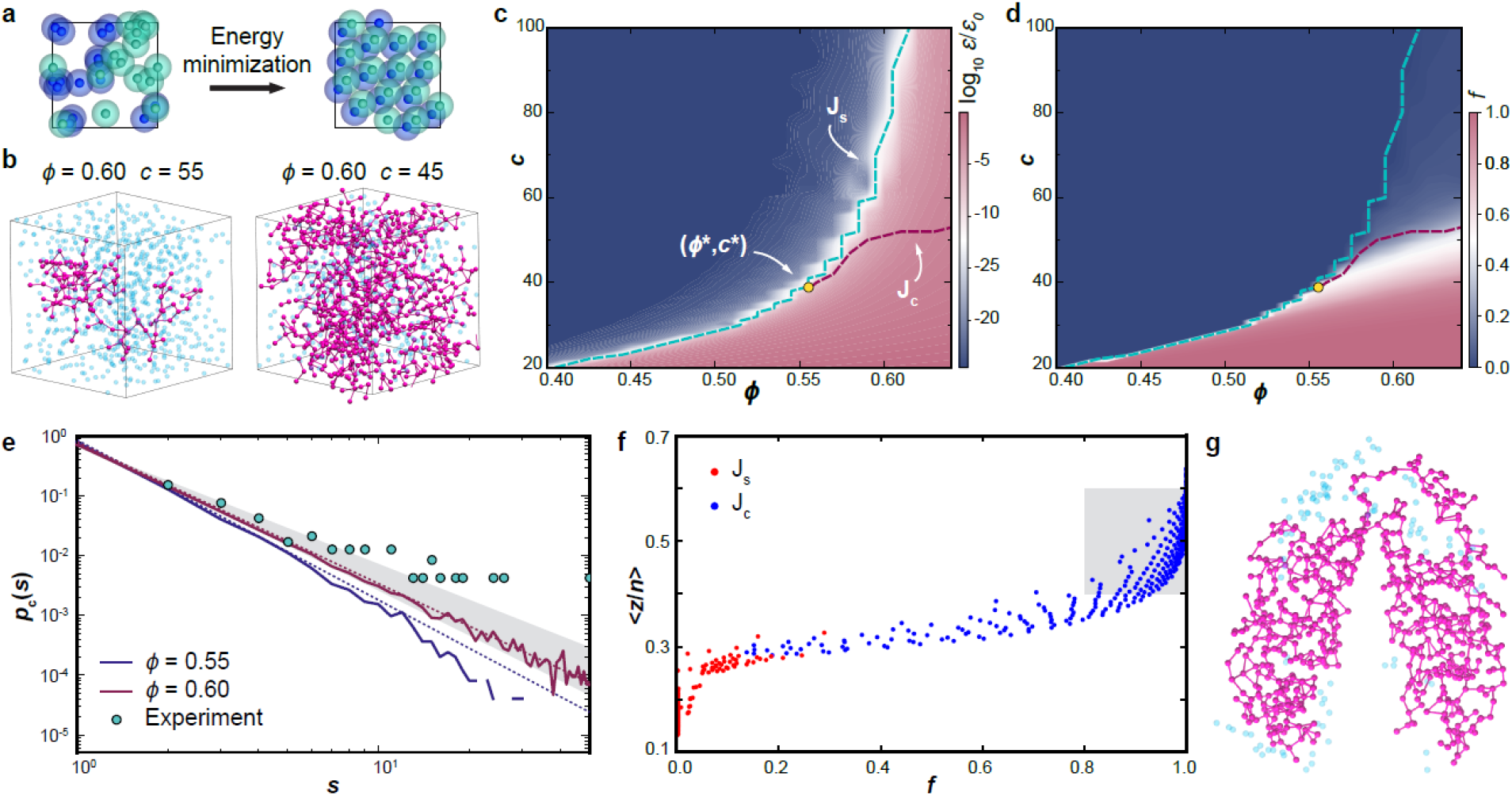
A chromatic jamming model for neuronal packing. **a**) Model schematic: blue and green particles represent distinct neuronal types, with a translucent zone around each representing the homotypic exclusion volume. **b**) Representative largest homotypic cluster (magenta) in chromatically unjammed (left) and jammed (right) states. Cyan: centroids of homotypic particles not in the largest cluster. **c**) Heatmap showing residual energy per particle after equilibration as a function of *c* and *ϕ*. Cyan line: J_s_ transition; red line: J_c_ transition. The triple point, (*ϕ*, c*\*), is localized at the intersection of J_s_ and J_c_. The range of *c* is set to match previous reports^13,26,27^ that the planarian brain contains several dozen neuronal types. **d**) Heatmap showing the fraction of particles belonging to the largest homotypic cluster **f**) as a function of *c* and ϕ. **e**) Homotypic cluster size distributions near the J_c_ transition exhibit power-law scaling. Solid line: simulation; dashed line: power-law asymptotics; symbols: experimental distribution, obtained by lumping all neuronal types to gain statistics. Note that distributions below the triple point, where *ϕ < ϕ*\*, diverge from the power law at large cluster sizes (e.g., *ϕ* = 0.55). Grey zone specifies the regime with scaling exponent (*τ*) between 2 and 2.5. **f**) Model predictions of 〈*z/n*〉, plotted against *f*, for simulations that cross the J_s_ (red) or J_c_ (blue) transition. Grey zone specifies the regime observed in experiments. **g**) Percolating homotypic cluster (magenta) detected in a *1020HH-1*^+^ neuronal population of the planarian brain. Cyan: centroids of *1020HH-1*^+^ neurons not in the percolating cluster.

**Fig. 3b** reveals an important signature of chromatic jamming: reducing *c* causes the homotypic contacts to cluster in space, even if the overall packing density *ϕ* remains unchanged. The sizes of these clusters grow until the largest homotypic contact cluster eventually spans the entire system. **Supplementary Fig. 4** shows that only under these conditions does the abundance of homotypic contacts reach values consistent with experiments. These observations suggest that the chromatic jamming transition may be analogous to the percolation threshold – at which long-range connectivity permeates a system – in random network models, which has been broadly used to describe gelation^24^.

To test this hypothesis, we sought to quantitatively define the transition point of chromatic jamming. We constructed a phase diagram by systematically varying *ϕ* and *c* across a biologically realistic regime. At low *ϕ*, the system is at the unjammed state and the energy per particle (*ε*) is practically zero (**Fig. 3c**). At sufficiently high packing density, a finite minimum energy indicates a spatial jamming transition, J_s_, at which all particle positions become constrained^20,21^ through both physical and homotypic contacts (**Supplementary Fig. 3b** and **Supplementary Fig. 5**). The critical density, *ϕ*_s_, increases with *c* and approaches the upper limit, the random close packing volume fraction in unicomponent systems^17–20^, 0.64, when *c* becomes sufficiently large. However, along the J_s_ line, the homotypic contribution to the system energy increases as *c* reduces (**Supplementary Fig. 5**), suggesting energy alone is insufficient for specifying chromatic jamming.

Next we explored the fraction of particles belonging to the largest homotypic cluster, *f*, which serves as the order parameter orthogonal to energy in a percolating system^24^ and characterizes the configurational degree of freedom in organizing cell types. **Fig. 3d** shows that the regime in which *f* approaches unity in our model deviates from the J_s_ line, especially at high packing density.

The basic ansatz of percolation^24^ states that the distribution *p*(*s*) of homotypic clusters of size *s* near the transition has a universal power law shape (see **Methods**). We identified conditions under which the power-law distribution persists over the widest range of *s*, as *p*(*s*) should deviate from the power law preceding and succeeding the transition (**Fig. 3e** and **Supplementary Fig. 6**). These conditions quantitatively specify the chromatic jamming transition, J_c_, representing the minimum number of types required to prevent chromatic jamming (**Fig. 3c-d**). Notably, unlike conventional jamming transitions, J_c_ plateaus at high packing density *ϕ*, where physical contacts in the system become rather invariant^17,18^. The intersection of the J_s_ and J_c_ lines gives rise to a triple point (*ϕ*, c*\*) = (0.56, 40). Below *ϕ*\*, J_s_ and J_c_ occur simultaneously, as the contribution of homotypic contacts dominates that of physical contacts (**Supplementary Fig. 5b**); above *ϕ*\*, J_s_ and J_c_ diverge and chromatic jamming emerges as a distinct jammed state. While the exact location of the triple point may be specific to model parameters, we anticipate that its existence is a generic feature of chromatic jamming.

These simulation results reveal the percolation nature of chromatic jamming and suggest several experimentally quantifiable hallmarks. First, the size distribution of homotypic neuronal clusters at the J_c_ transition should follow a power law for 3D percolation^24^, with the exponent *τ* ≈ 2.2, which is nearly constant along the J_c_ line when *ϕ* > *ϕ*\* (**Supplementary Fig. 6**). This is exactly as observed in the experiment, albeit finite-size effects – due to relatively small numbers of homotypic neurons in individual brains – limit the statistics of larger clusters and cause a deviation at the tail (**Fig. 3e**). Second, experimentally observed 〈*z/n*〉 ≈ 0.4-0.6 for homotypic neuronal neighbors should only exist after the J_c_ transition (**Fig. 3f** and **Supplementary Fig. 4**). This is distinct from the conventional jamming model in which 〈*z/n*〉 is a function of packing density^17–19^. Third, large percolating homotypic clusters should be detectable at the J_c_ transition and are accordingly found in the planarian brain, as exemplified in **Fig. 3g.** These lines of evidence collectively indicate that the neurons in the planarian brain indeed pack at the J_c_ transition state.

The chromatic jamming transition described here extends the classic understanding of jamming by introducing cell diversity as an orthogonal dimension, beyond the traditional physical constraints governing analogous transitions in both living and inanimate matter^1,3,4,6,17,18,20^. Cell type-specific interactions impose additional geometric constraints that favor a subset of packing configurations, resulting in a previously uncharacterized jammed state. Hallmarks of such a state have been quantitatively verified in the brain of a model organism, the planarian. Although described in a basal nervous system, chromatic jamming may also provide a general physical framework for neurons to organize in other densely packed neural tissues. The structural constraints in chromatic jamming can have important physiological implications in a variety of biological processes, including neuronal differentiation, cell type diversification, and optimization of neuronal network connection efficiency. More broadly, this work reveals a living example of a complex network structure with multiple percolated sub-networks intercalating in space, which may find other applications, as in signal communication and graph optimization.

## Methods

### Animals

Asexual *S. mediterranea* (CIW4 strain) were maintained in the dark at 20°C in ultrapure water supplemented with 0.5 g/L Instant Ocean Sea Salts and 0.1 g/L sodium bicarbonate. Planarians were fed calf liver paste once or twice weekly and starved at least 5 d prior to all experiments.

Most imaging was performed on wide-type planarians, but to quantify the spatial arrangement of neurons in ectopic neural tissues (**Supplementary Fig. 2**), *β1-integrin* knock-down planarians were obtained with feeding of double-stranded RNA (dsRNA) to induce RNA interference (RNAi) against the dd_Smed_v6_2017_0_1 transcript (PlanMine), following the procedure described in Refs. 29 and 30. dsRNA was synthesized by *in vitro* transcription from the clone generated using oligonucleotide primers (5’ to 3’) GAACTCAACACACAACGCCC and TCTCGACAGGGAACAATGGC to amplify the gene fragment from cDNA and clone into vector pJC53.2 (Addgene Plasmid ID: 26536)^28^. dsRNA was mixed with liver paste at a concentration of 100 ng/μL and fed to planarians 3 times in 5 d. Planarians were head amputated 4 hr after the final feeding and allowed to regenerate 16 d before fixation.

### RNA-FISH and image analysis

RNA probes against neuropeptide genes were synthesized as described previously, and FISH was performed following the established protocol^28,31^. To determine centroids of individual neurons using FISH signals, confocal stacks were obtained from a laser-scanning confocal microscope (Zeiss) at over-sampled resolutions, i.e., 350 nm in lateral and 700 nm in axial using a 20x N.A. = 0.8 objective, as recommended by Imaris (Bitplane). The stacks were then resampled to give isotropic voxels and subjected to Gaussian filtering and background subtraction. Centroids of labeled cell bodies were segmented channel-by-channel with Imaris using parameters empirically determined to minimize the need for manual curation. The average cell size was determined to be 6 μm, which also sets the upper limit for the fitting error in determining centroids. The centroid positions in 3D are then extracted for downstream statistical analyses.

Voronoi tessellation was performed on centroids using Delaunay triangulation in MATLAB. To define nearest neighbors that are also in “contact” for each individual neuron, we first identified the closest nearest neighbor, then counted the number of nearest neighbors that are the same distance away within an error tolerance range, which is the centroid fitting uncertainty, 6 μm. Granocentric model was plotted using the model and parameters specified previously^17,19^, with inverse Laplace transform computed numerically.

### Simulation and statistical analysis

Cells are represented as soft particles of uniform size interacting with a Hookean potential of the form *u*(*r*) = 0.5*ϵ*(1−r/*σ*)^2^ for *r* ≤ *σ*, and 0 otherwise, where *u*(*r*) is the pairwise potential between two particles separated by distance *r*, and *σ* and *ϵ* are the range and strength of interaction, respectively. Every particle is randomly assigned a color. The interaction range and strength for pairs of heterotypic particles are *σ* = *σ*_0_ and *ϵ* = *ε*_0_, whereas those for homotypic, concolor particles are *σ* = 3*σ*_0_ and *ϵ* = 3*ε*_0_.

Every system contains *N* particles divided evenly into *c* types. Multiple system sizes were explored and the results reported are obtained from those performed on the largest system containing *N* = 40,000 particles to minimize size dependence. *N* was adjusted manually in each simulation to ensure that the number of particles for each type is identical. The simulation box is cubic and its dimension is determined from the particle packing density through *ϕ* = *Nπσ*_0_^3^/(6*V*), which gives a box edge greater than 30*σ*_0_ for all densities studied. Energy minimization was carried out in LAMMPS^32^ until the energy change per particle between consecutive simulation steps plateaus or reaches a tolerance of 10^−20^ *ε*_0_ (**Supplementary Fig. 3**). Steep descent was used first to relax the system to a configuration near the local minimum, followed by a Hessian-free truncated Newton algorithm to further minimize system energy. The periodic boundary condition was imposed.

Three independent configurations were simulated for each condition and the energy per particle (*ε*) was computed by averaging across all configurations. The J_s_ point was then identified by the steepest jump of the energy, typically across several orders of magnitude, as varying packing density or color number (**Supplementary Fig. 3**). The J_c_ point was identified following the basic ansatz of percolation^24^, which states that *p*(*s*) near the transition has a universal shape, *p*(*s*) = *s*^−*τ*^ *g*(*s|c* − *c*_h_|^1/*σ*^), in which *g* is a cutoff function that plateaus for small *s* and damps rapidly for large *s*. The term |*c* − *c*_h_|^1/*σ*^ is the reciprocal of the typical cluster size, which vanishes at ch. Geometric analysis was performed using Voronoi tessellation, similarly to the experimental protocol. For cluster analysis, two particles separated by a distance shorter than a threshold *l_c_* were assigned to the same cluster. A value *l_c_* = 2.99995*σ*_0_ was used to account for the homotypic repulsion with a truncation error. The transition point *c*_h_ is then located at the curve exhibiting the power-law distribution over the widest range of *s*, as *p*(*s*) should deviate from the power law preceding and succeeding the transition (**Supplementary Fig. 5**).

### Code availability

Analysis and simulation codes are available for public access on GitHub (https://github.com/xianshine/cJamming).

### Data availability

The datasets generated and analyzed within this study can be downloaded from https://github.com/xianshine/cJamming or available from the corresponding authors upon request.

## Supporting information

## Acknowledgements

We thank Kejia Chen and Ke Chen for help with data analysis, R. H. Roberts-Galbraith and J. J. Collins III for experimental assistance during the early phase of this study, S. Granick, I. Riedel-Kruse, D. M. Sussman, and P. N. Newmark for discussions. Plasmids that contain the planarian neuropeptide genes are kindly provided by J. J. Collins III. This work is supported by the Burroughs Wellcome Fund through the CASI program to B.W.

## Author contributions

J.Q. and B.W. designed the research, M.K. and B.W. performed the experiments, M.K., X.K. and B.W. analyzed the data, X.K. performed the simulation, X.K. and J.Q. analyzed the simulation results, all authors wrote the paper.

