## Supplementary material for "‘Chromatic’ Neuronal Jamming in a Primitive Brain"

**Supplementary Figure 1-6.**

**Supplementary Note 1.**

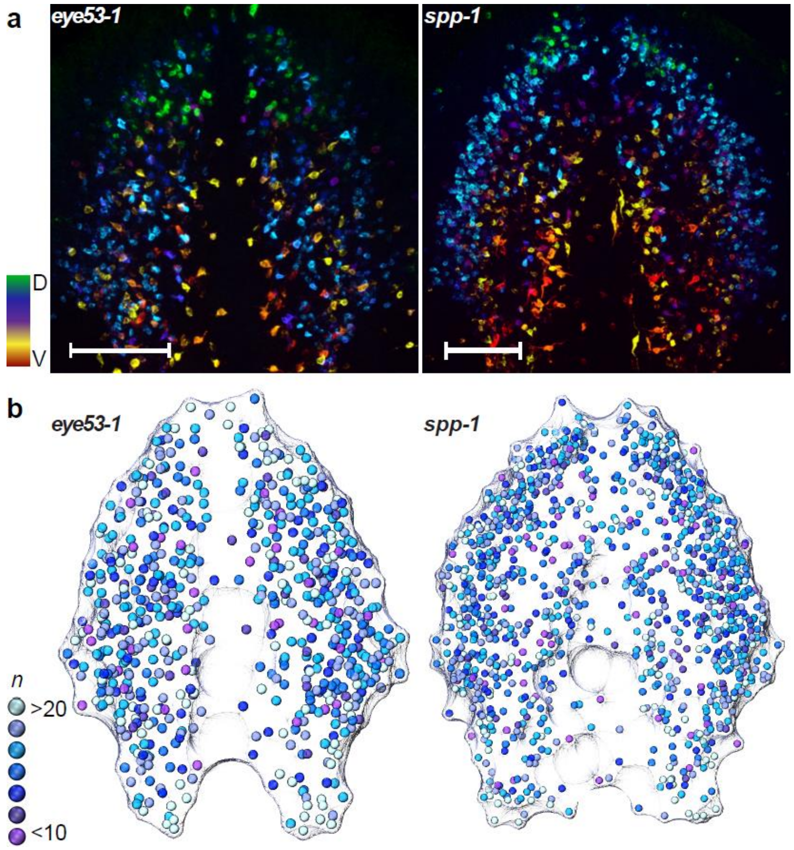


**Supplementary Figure 1. Single-neuron resolution of multiple peptidergic neuronal types in the planarian brain. a**) Depth-colored projection images showing RNA-FISH of neuropeptides *eye53-1* (left) and *spp-1* (right) to reveal spatial distributions of *eye53-1*$^+$ and *spp-1*$^+$ peptidergic neurons, respectively, in the planarian brain. Colormap: dorsal-to-ventral. Scale bars: 100 μm. **b**) Centroids of *eye53-1*$^+$ (left) and *spp-1*$^+$ (right) peptidergic neurons corresponding to images in **a**. Colors represent number of nearest homotypic neighbors (*n*) of each individual neuron, determined by Voronoi tessellation.

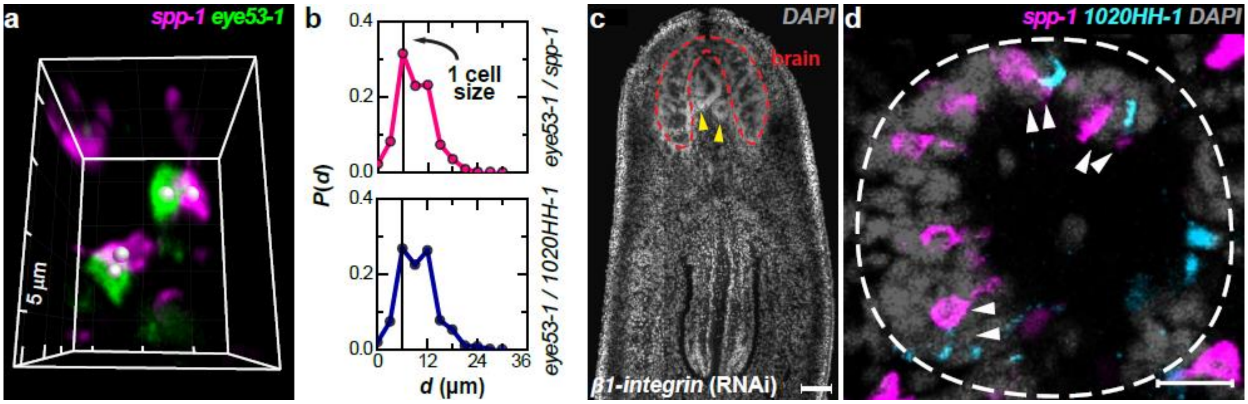


**Supplementary Figure 2. Repulsion is not observed between heterotypic neuronal types. a**) 3D reconstruction image showing that *spp-1*$^+$ (magenta) and *eye53-1*$^+$ (green) neurons can locate in physical proximity. Grey: cell body centroids. **b**) Probability distributions of separation between closest pairs of heterotypic neurons for 3 representative peptidergic neuronal types: *eye53-1*$^+$ / *spp-1*$^+$ (top) and *eye53-1*$^+$ / *1020HH-1*$^+$ (bottom). Statics on heterotypic pairs were obtained by analyzing 4 planarians each. Distributions peak around 6 μm, equivalent to one cell size, suggesting heterotypic neurons often form physical contacts. **c**) To confirm that inter-neuronal spacing is controlled by local cell-cell interactions rather than global patterning cues that may be specific to the brain tissue, neuronal organization is characterized in ectopic neuronal aggregates (yellow arrowheads). Ectopic neural spheroids are induced by knockdown of *β1-integrin* using RNA interference (RNAi) as reported previously[29,30]. Scale bar: 100 μm. **d)** A representative spheroid (dashed circle) is shown to illustrate that heterotypic neurons (*spp-1*$^+$ and *1020HH-1*$^+$ peptidergic neurons) can locate in physical proximity to form dimers (white arrowheads), whereas homotypic neurons are always separate in space. This data is consistent with homotypic repulsion, even in this ectopically induced neural mass outside of the brain. Scale bar: 20 μm.

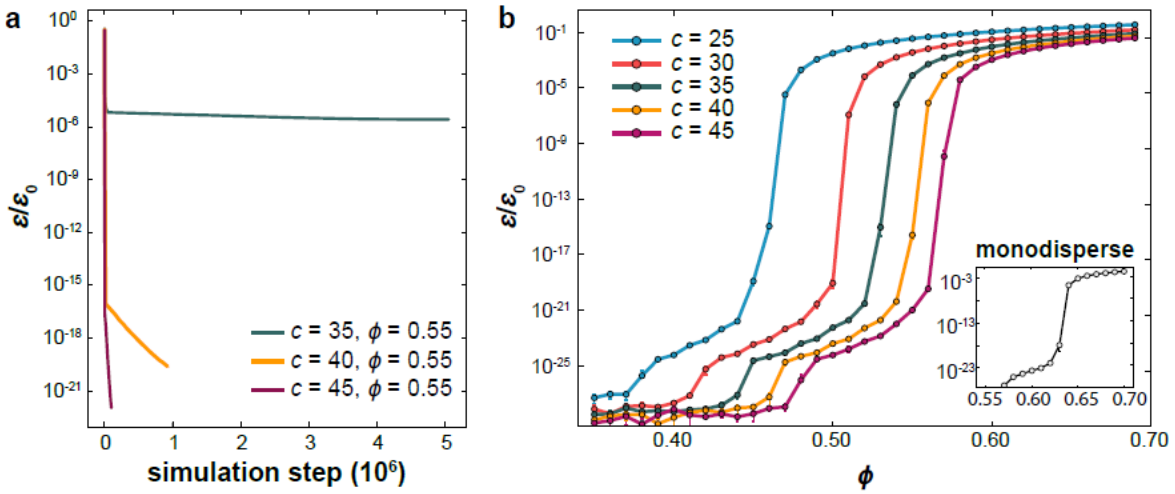


**Supplementary Figure 3. System energy defines the $J_s$ transition. a)** Residual energy per particle ($\varepsilon$) evolution during the energy minimization equilibration for several color numbers ($c$) at a fixed packing fraction ($\phi$). Simulations were terminated when energy change between consecutive steps drops below a tolerance of $10^{-20}\,\varepsilon_0$. **b)** Minimum residual energy per particle ($\varepsilon$) versus $\phi$ for varying $c$. The $J_s$ transition is defined at critical $\phi$ when the steepest increase in energy of each condition is identified. Inset: energy residual varying with $\phi$ in a monodisperse system. The steepest energy jump occurs near the well-defined jamming point[17,19] of $\phi_s = 0.64$. Error bars represent standard deviation.

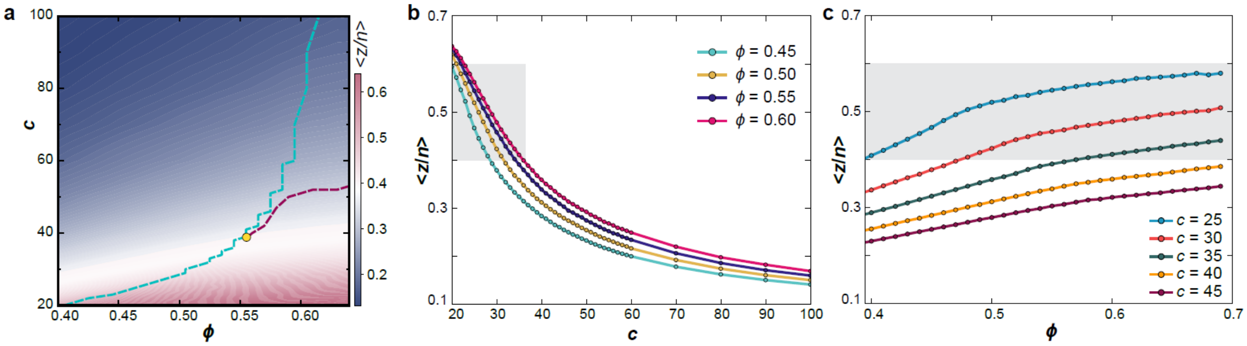


**Supplementary Figure 4. The average fraction of contact neighbors, ⟨*z*/*n*⟩, of homotypic neighbors depends strongly on color number but weakly on packing density. a)** Heatmap depicting ⟨*z*/*n*⟩ of the model as a function of *c* and $\phi$. Cyan line: $J_s$ transition; red line: $J_c$ transition. **b**) Simulated ⟨*z*/*n*⟩ is plotted against *c* for varying $\phi$. **c**) ⟨*z*/*n*⟩ is plotted against $\phi$ for varying *c*. Note that, in conventional jamming models, ⟨*z*/*n*⟩ is expected to be a function of packing density[17-19]. Grey zones specify the regime observed in experiments.

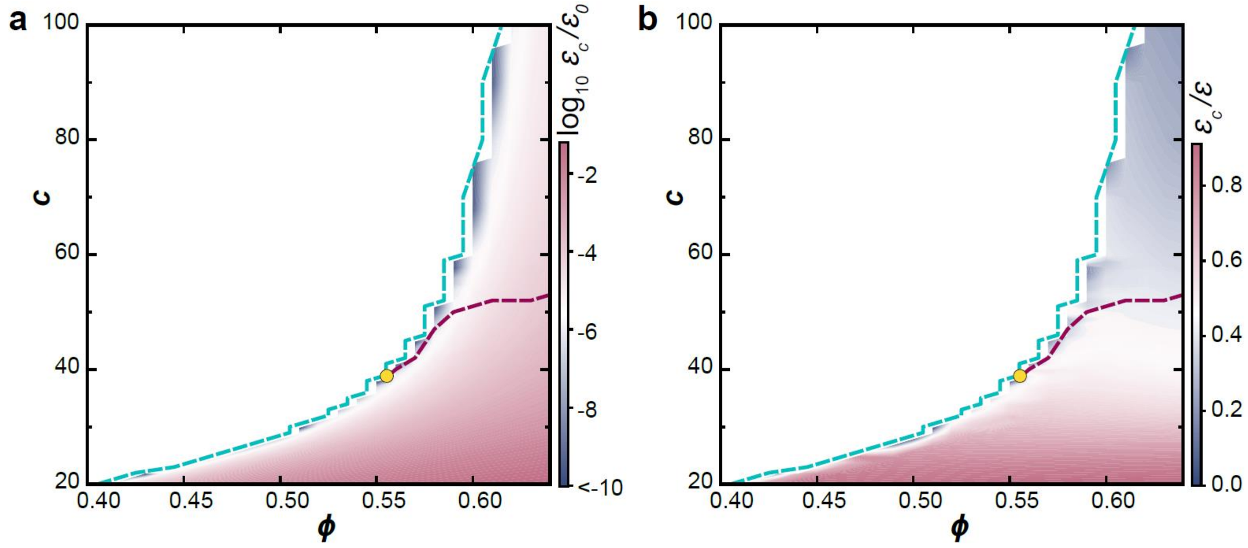


**Supplementary Figure 5. Energy contribution of homotypic interactions increases as color number decreases. a)** Heatmap showing the residual energy per particle due to homotypic interactions as a function of $c$ and $\phi$. **b)** Heatmap depicting the fraction of total energy that arises from homotypic interactions, as a function of $c$ and $\phi$. Cyan line: $J_s$ transition; red line: $J_c$ transition.

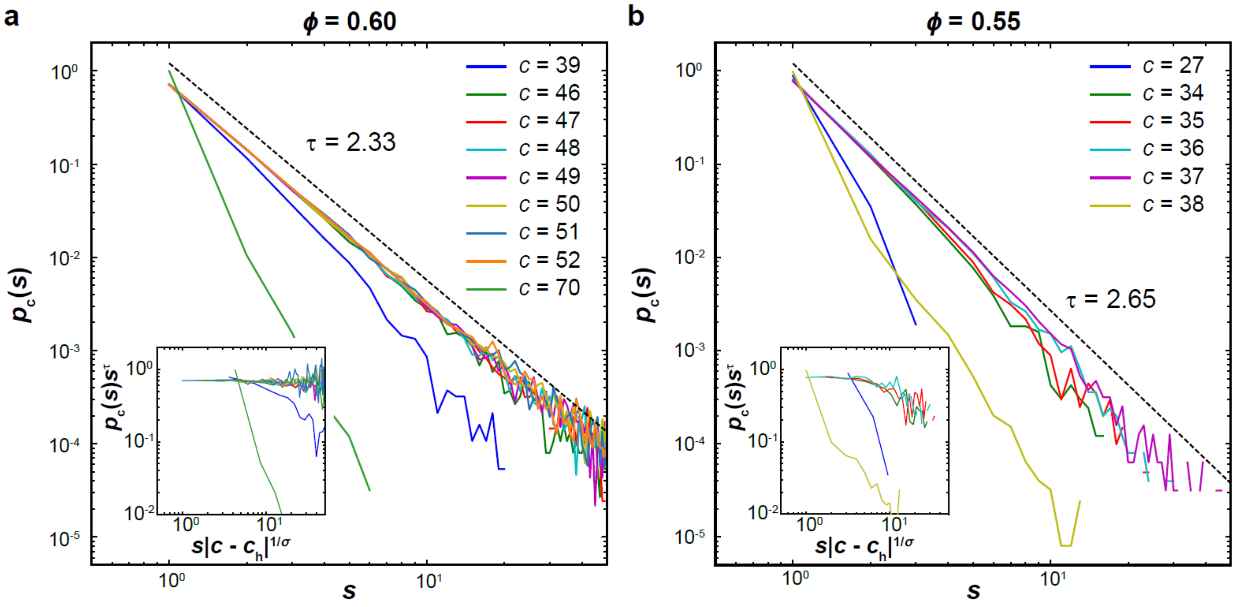


**Supplementary Figure 6. Power-law cluster size distribution defines the $J_c$ transition.** Cluster size distributions at two packing densities, **a**) $\phi = 0.60$, and **b**) $\phi = 0.55$, flanking the triple point ($\phi^*$, $c^*$) = (0.56, 40). Distributions with several color numbers near the critical color number ($c_h$) are shown; the $J_c$ transition is defined when the power-law distribution spans the widest range of *s*. For comparison, two cases far away from $c_h$ are shown as divergent from the power law. Dashed lines: asymptotic power law with exponent $\tau$ specified. $\tau$ is consistent with 3D percolation theory[24]. Inset: the collapse of cluster distributions as $p_c(s)s^\tau$ is plotted against $s|c-c_h|^{1/\sigma}$ with corresponding $\tau$ and $\sigma = 2$. Note that, when $\phi > \phi^*$, the power-law distribution is robust throughout broad ranges of color numbers around $c_h$, whereas the power law only exists in narrower ranges as $\phi < \phi^*$ and bends down at large cluster sizes. Moreover, τ remains nearly constant when $\phi > \phi^*$. Below $\phi^*$, even though an analogous exponent τ can be still determined for finite clusters, it grows steadily as $\phi$ decreases. Why τ deviates from the classic value below $\phi^*$ remains a subject of future study.

### Supplementary Note 1: Homotypic repulsion

It is self-evident that the distribution of neurons in any neural tissue must be tightly regulated through molecular cues specifying the identities of cells that surround any given cell[9,10]. These cell type-specific interactions can be mediated by diffusive factors that are recognized by cells at a distance or through cell surface receptors distributed along extended cellular processes. Indeed, diverse molecular mechanisms may be involved, as shown across a variety of nervous systems, including those of *Drosophila*, mouse, and *C. elegans* (e.g., transmembrane cadherin Flamingo, TGF-b/Activin, immunoglobulin superfamily cell adhesion molecules (IgCAMs), and integrin, among others)[10,11,23]. However, the exact molecular method by which homotypic interactions occur does not affect the characterization and understanding of neuronal packing, as it is an emergent problem that exists on a length scale much larger than specific molecular interactions, as supported by previous work in the field[8,9,22].

The precise molecular mechanism controlling the homotypic repulsion among planarian neurons remains to be uncovered, although many key molecular players of homotypic interactions (e.g., protocadherins and IgCAMs) are known to be deeply conserved in the planarian[33]. For example, neuronal spacing in the fly and mouse visual systems is known to be mediated by homotypic repulsion through Down syndrome cell adhesion molecule (DSCAM), a member of the IgCAM superfamily[11,22]. Homologs of IgCAMs, including DSCAM and neural cell adhesion molecule (NCAM), are known to be present in the planarian central nervous system, with functional analyses showing them as playing key roles in neuronal interactions[33]. Our experiments also suggest that DSCAM contributes to the homotypic repulsion between neurons in the planarian brain, but it is likely that DSCAM functions only as one of several parallel or redundant mechanisms. Precisely defining the molecular mechanism underlying neuronal homotypic repulsion requires extensive genetic studies coupled with quantitative measurements of spatial neuronal organization in the planarian brain, a methodology this study aims to establish. Nonetheless, such a mechanism is not required to understand the generic physical principles underlying the cellular packing on a global level, and therefore is beyond the scope of this study.
